# Depositing biological segmentation datasets FAIRly

**DOI:** 10.1101/2024.12.10.627814

**Authors:** Elaine ML Ho, Dimitrios Ladakis, Mark Basham, Michele C Darrow

**Author notes:** Corresponding author: Michele Darrow.

## Abstract

Segmentation of biological images identifies regions of an image which correspond to specific features of interest, which can be analysed quantitatively to answer biological questions. This task has long been a barrier to conducting large-scale biological imaging studies as it is time- and labour-intensive. Modern artificial intelligence segmentation tools can automate this process, but require high quality segmentation data for training, which is challenging to acquire. Biological segmentation data has been produced for many years, but this data is not often reused to develop new tools as it is hard to find, access, and use. Recent disparate efforts (Iudin, et al., 2023; Xu, et al., 2021; Vogelstein, et al., 2018; Ermel, et al., 2024) have been made to facilitate deposition and re-use of these valuable datasets, but more work is needed to increase re-usability. In this work, we review the current state of publicly available annotation and segmentation datasets and make specific recommendations to increase re-usability following FAIR (findable, accessible, interoperable, re-usable) principles (Wilkinson, et al., 2016) for the future.

## Introduction

Segmentation is a required process for many volumetric biomedical imaging experiments and other bioimaging studies. It is the step where scientists derive either illustrative examples or quantitative information which allows them to address the scientific question(s) at hand. For example, segmentation has been used to understand many aspects of biomedical health research and basic science research, from the cellular composition of the blood-nerve barrier (Malong, et al., 2023), the structure of the vascular network in equine placenta (Laundon, et al., 2024), the effects of SARS-CoV-2 viral infection (Mendonça, et al., 2021), and even as a tool for development of novel imaging techniques (Dumoux, et al., 2023). Across medical and basic science research, and imaging modalities from electrons (vEM, SEM, TEM) to X-rays (SXT, microCT), segmentation is a crucial step.

Until recently, segmentation was completed manually by tracing the outlines of the features of interest on all or many slices through a 3D dataset. With the advent of machine learning, this task can now be semi-automated using shallow (Luengo, et al., 2017) and deep learning approaches (Belevich, Joensuu, Kumar, Vihinen, & Jokitalo, 2016; Conrad & Narayan, 2023; Pennington, et al., 2022; Arganda-Carreras, et al., 2017; Berg, et al., 2019; Thermo Fisher Scientific) using foundational models (Conrad & Narayan, 2023; Narayan & Conrad, 2022). Even with these advances, segmentation still requires varying amounts of manual annotation to provide training data for machine learning approaches to learn from. In most cases, machine learning segmentations also require some level of manual correction to provide a high-quality output that can be used to derive meaning. Because of this level of manual intervention, high-quality segmentations are a scarce and valuable resource (Czymmek, et al., 2024). However, as with many scientific endeavours, the details associated with how the segmentations are produced are vital to downstream re-use. Our goal was to understand the current state of electron and X-ray image segmentation deposition and the possibilities and logistics associated with reuse.

There are a small number of options for public deposition of segmentations. The standard option in the vEM and X-ray imaging fields is EMPIAR (Electron Microscopy Public Image Archive) (Iudin, et al., 2023) which was created in 2013 to address the need for a raw/image data repository in the fields of cryo-electron microscopy and tomography (cryoEM and cryoET), though it has now expanded its remit to include vEM and X-ray imaging. As of writing, EMPIAR has released just shy of 5 PB of data across >1,800 entries. While it has the distinction of being the first community repository for segmentation data, its utility has so far been limited by the lack of standardised metadata collection during deposition and its broad purview. Prior to EMPIAR, the Electron Microscopy Data Bank (EMDB) (The wwPDB Consortium, 2024) acted as a repository for electron microscopy output maps, including in a small number of cases, segmentation maps uploaded as accompanying datasets. Smaller, more bespoke repositories have been created for specific purposes. For example, OpenOrganelle (Xu, et al., 2021; Heinrich, et al., 2021) was created to host the vEM datasets and accompanying segmentations specifically associated with the enhanced focused ion beam SEM (eFIB-SEM) technology that was developed at HHMI Janelia (Xu, et al., 2017). Similarly, some research groups with a novel technique (nanotomy.org; (Pirozzi, Hoogenboom, & Giepmans, 2018)) have generated their own open repositories for storage of data and associated metadata. Some segmentation data and metadata can also be found on open generic repositories like Zenodo (European Organization For Nuclear Research & OpenAIRE, 2013) or associated with public challenges such as on Kaggle (Kaggle, n.d.). Finally, most recently, the Chan Zuckerberg Imaging Institute has entered this area with the CryoET Data Portal (Ermel, et al., 2024), which at time of writing, contains ∼300 entries (datasets) mainly focused on cryoET data with annotated particle locations. 14 of these depositions have ”ground truth” annotations, nine of which are also found on EMPIAR. Importantly, the depositions have a standardised metadata structure, naming conventions, application programming interface (API), and detailed curation of metadata.

Here, we undertook a landscape review of the publicly available segmentation data and associated metadata to understand and report on gaps and barriers within the current system, as well as recommendations to address barriers to re-use.

## Results

Datasets associated with electron and X-ray imaging segmentations (image, training, and label data) were searched for in common repositories which are known to contain segmentation data from these techniques (e.g., EMPIAR, EMDB, OpenOrganelle). Other general microscopy repositories such as the BioImage Archive (Hartley, et al., 2022) or the Image Data Resource (Williams, et al., 2017) were not specifically searched due to their inclusion and focus on other imaging modalities which are outside the scope of this work. Publications associated with segmentation were searched for using PubMed and a representative sub-sample was selected (see Methods). Searches were limited to publications from 2014 – 2024, and across all searches a total of 227 unique publications were selected. These publications were then assessed for various characteristics associated with segmentation datasets, metadata and re-use. These characteristics included:

- Type of study: biological (answering a biological question), method (sample preparation or image acquisition), software (software tool for processing the image data), dataset (describing a useful dataset for the community).
- Purpose of segmentation: Qualitative visualisation, quantitative morphology, proof-of-concept, benchmarking, single particle analysis (SPA) or sub-tomogram averaging (STA).
- Repository where data was deposited to, if at all. Here, we recorded the deposition location of four categories of data: the source data (microscopy images upon which segmentations are based), training data (microscopy images and their labels which were used to train artificial intelligence models), label data (segmentation masks), and code.
- Imaging modality
- File formats for source, training, and label data
- Origin of data, i.e. whether this publication generated new source/raw data or segmentations, and if they were generated elsewhere, where did this data come from
- Segmentation type: semantic (categorises all objects of a class as one, e.g. all mitochondria have one value in the segmentation mask) or instance (each object in a class is separated, e.g., individual mitochondria have different values in the segmentation mask)
- Segmentation method: manual, semi-automated, automated
- Biological scale and/or tissue

Based on this landscape analysis, we have identified both characteristics associated with data reuse and barriers preventing reuse of published segmentation data according to FAIR principles – scientific data should be findable, accessible, interoperable, and reusable (Wilkinson, et al., 2016).

### Findability and accessibility of deposited data

Data cannot be reused unless the data and its associated metadata can be found and accessed. Of the 227 publications surveyed, 64.4% of source data, 76.3% of training data, 82.5% of label data, and 55.6% of code was either not deposited, available only on request, or not available anymore (**Error! Reference source not found.**). In some cases, data was previously deposited in locations that were no longer available, e.g., at websites that no longer exist. Some data was deposited in locations that were accessible, but difficult to find. For example, the reader would need to search through citation chains or using keywords online to find the correct dataset. Amongst the findable and accessible locations, the most common location for source data and label data to be deposited was in field-specific databases such as EMPIAR (Iudin, et al., 2023), EMDB (The wwPDB Consortium, 2024), or Neurodata (Vogelstein, et al., 2018). Another popular location is general databases such as Dryad (Dryad, n.d.) or Zenodo (European Organization For Nuclear Research & OpenAIRE, 2013); which also function similarly to field-specific databases, but due to their wide scope are much more difficult to search unless already aware of the existence of a specific entry.

Label data, i.e., the segmentation masks that were generated in the publication, was findable and accessible in 25% of the 223 publications surveyed. Quantitative measurements are often taken from the label data to draw biological conclusions, but the label data was not made available in 74 of the 99 publications that did this.

Reproducibility of scientific studies is important to ensure the integrity of the biological conclusions that we draw from our research. This is especially important as we move towards artificial intelligence approaches which are not fully explainable – in these cases it is crucial that the data and procedures for training models are made available for scrutiny to avoid bias. Unfortunately, this was not the case for many of the publications that were surveyed. Of the 97 publications surveyed which required a model to be trained, training data was unavailable or inaccessible for 74 of them (76.3%). Code for generating semi- or fully automated segmentations was available in 48.9% of publications surveyed. Note that in this study, code was considered “deposited” if there was a working link to the scripts or software used to generate the result, no tests were done to check if the results were reproducible from the code that was available.

The analysis for Figure 1 was repeated using only the 194 publications from the PubMed literature search to avoid possible over-representation of the field-specific databases that were also included in the wider results presented above (EMPIAR, EMDB, Open Organelle). As expected, the proportion of source, training, and label data that was not deposited at all was higher in this subset of publications that did not include the field-specific searches (Supplemental Figure ). Field-specific databases remained the most popular findable and accessible deposition location for source data (13.9%) but there was a more even split between public challenges (10.3%), self-maintained websites (8.8%), and general scientific databases (6.7%). Label data generated for public challenges is often saved as an entry to the public challenge, and therefore not often deposited elsewhere. The publications which did deposit label data were relatively evenly distributed between field-specific databases (5.7%), general scientific databases (4.7%) and self-maintained websites (5.2%).

**Figure 1:**
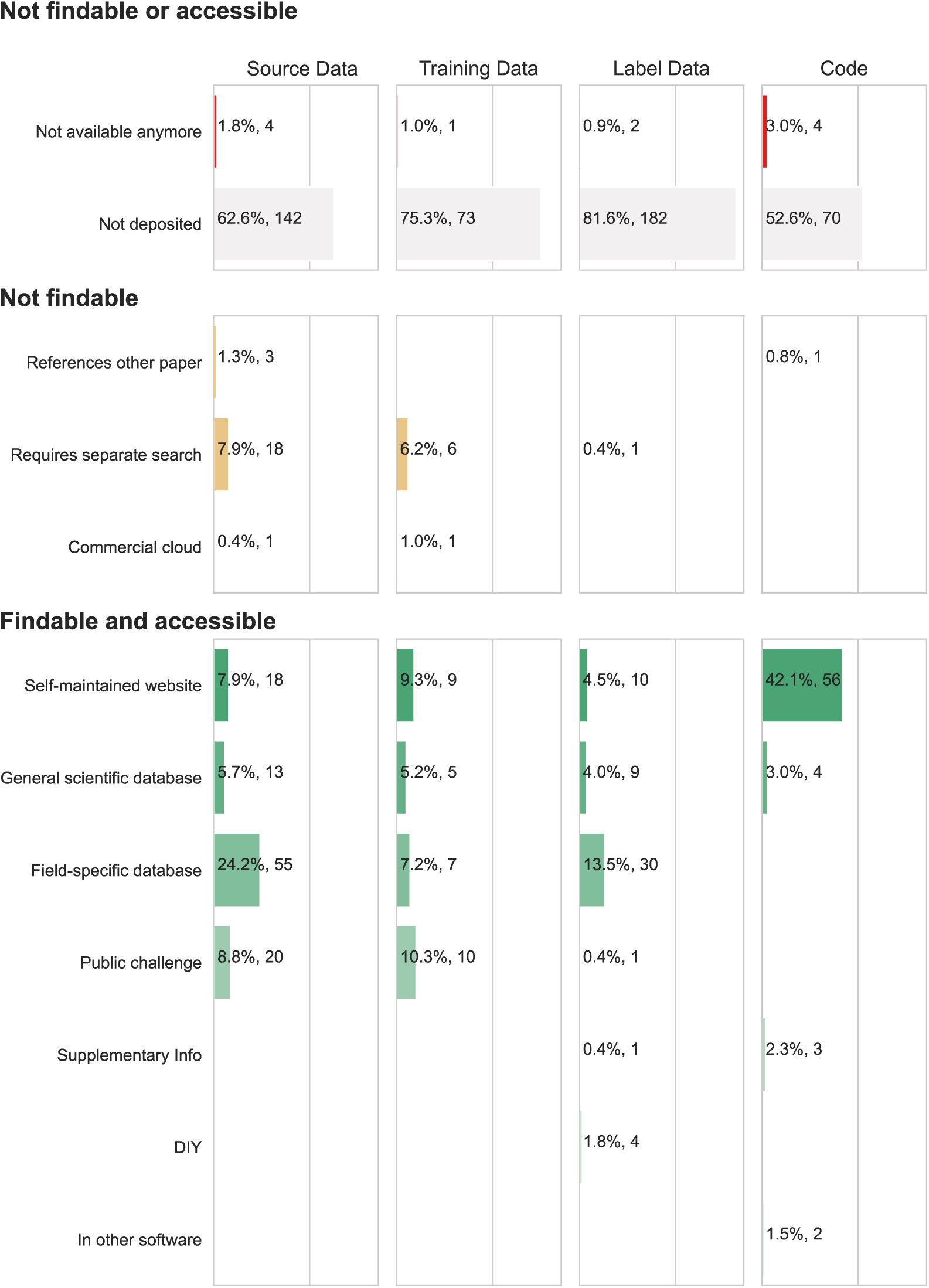
Deposition locations of data from publications surveyed (n=227). Most publications did not deposit their data at all, or deposited in locations which were no longer available, e.g., at defunct institutional repositories. Some of the depositions were accessible but diDicult to find, e.g., reader would be required to search for the data separately if the dataset was named but no direct working link was provided. Image data and label data was most likely to be deposited in field-specific databases such as EMPIAR or EMDB, whilst training data (where available) was most commonly available from public challenges. Code was most frequently available from self-maintained websites such as Github repositories. Note that some publications can be counted in multiple categories, e.g., if the publication had training data available online in a public challenge but required a separate search.

**Figure 2.**
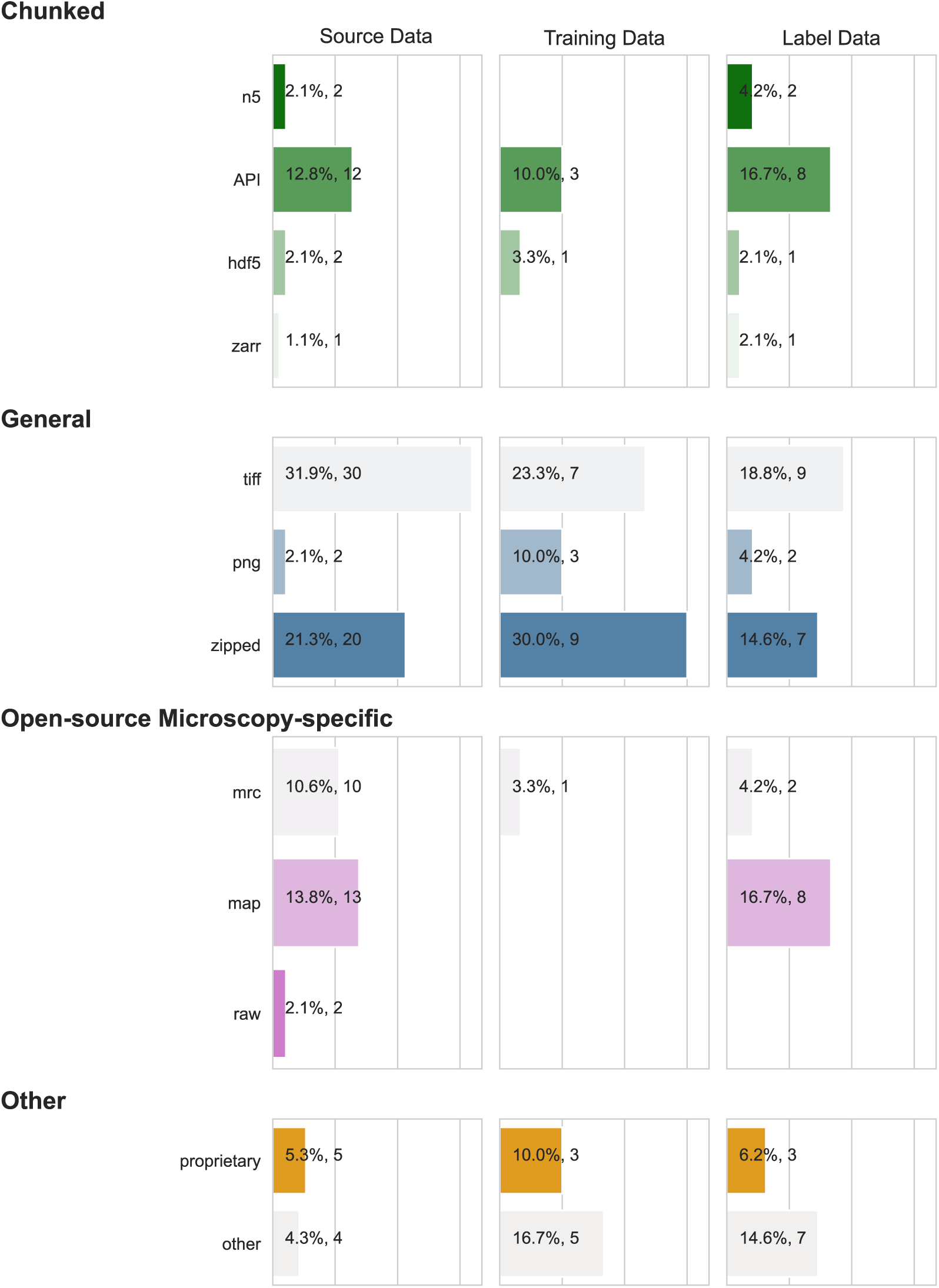
File formats for source, training, and label data for the surveyed publications (n=227). Note that some publications used multiple file formats for each category and are counted multiple times in this instance. Source, training, and label data was deposited in 26 diDerent file formats. Chunked or pyramidal file formats allow data to be accessed at diDerent resolution levels or region-of-interest sizes easily. Data accessed via an API are typically stored in a chunked format on cloud providers for easy programmatic access. General data formats are often used, with tiD images being the most popular across all data types. Many datasets were also uploaded as zipped files, so the file formats of the actual data were not easily known before downloading the data. Microscopy-specific file formats were quite common, especially the mrc and map formats (which are the same mrcfile format) which together made up 22.3% of image data, 3.1% of training data, and 20% of label data. Proprietary and other more obscure file formats were more common in training and label datasets than in source data.

### Interoperability of deposited data

Integrating data collected from different sources is also challenging due to the variability in file formats and metadata. Source, training, and label data was deposited in 26 different file formats (**Error! Reference source not found.**). This variability makes tasks such as combining similar segmentation datasets together more difficult, as customised code is often needed to parse this data into a common format. While many image processing software packages do accept data in different open-source formats, and tools for parsing and converting bioimaging file formats such as Bioformats (Linkert, et al., 2010) exist, this would still be a daunting task. This indicates that the lack of standardisation in file formats still presents a barrier to interoperability of segmentation data.

Metadata was not standardised across the various repositories, creating conflicts when merging data across repositories. In its most extreme cases, the definitions of some types of segmentation data can vary between fields, which could cause datasets to be used inappropriately by researchers unfamiliar with the nuances of different fields (see Discussion). In many cases, metadata was simply missing, making re-use nearly impossible without extensive field-specific knowledge.

### Rates of reuse

Rates of data reuse in biological segmentation studies are low. In all publications surveyed, only 23.3% did not generate new data for their study (Figure 3). Software publications collected new data for 57.7% of the studies, which is surprisingly high considering that most software is developed for established use cases, for which segmentation data likely already exists. Data reuse is more common in connectomics studies, with a slim majority of studies reusing data rather than collecting new data (55% reused data).

**Figure 3.**
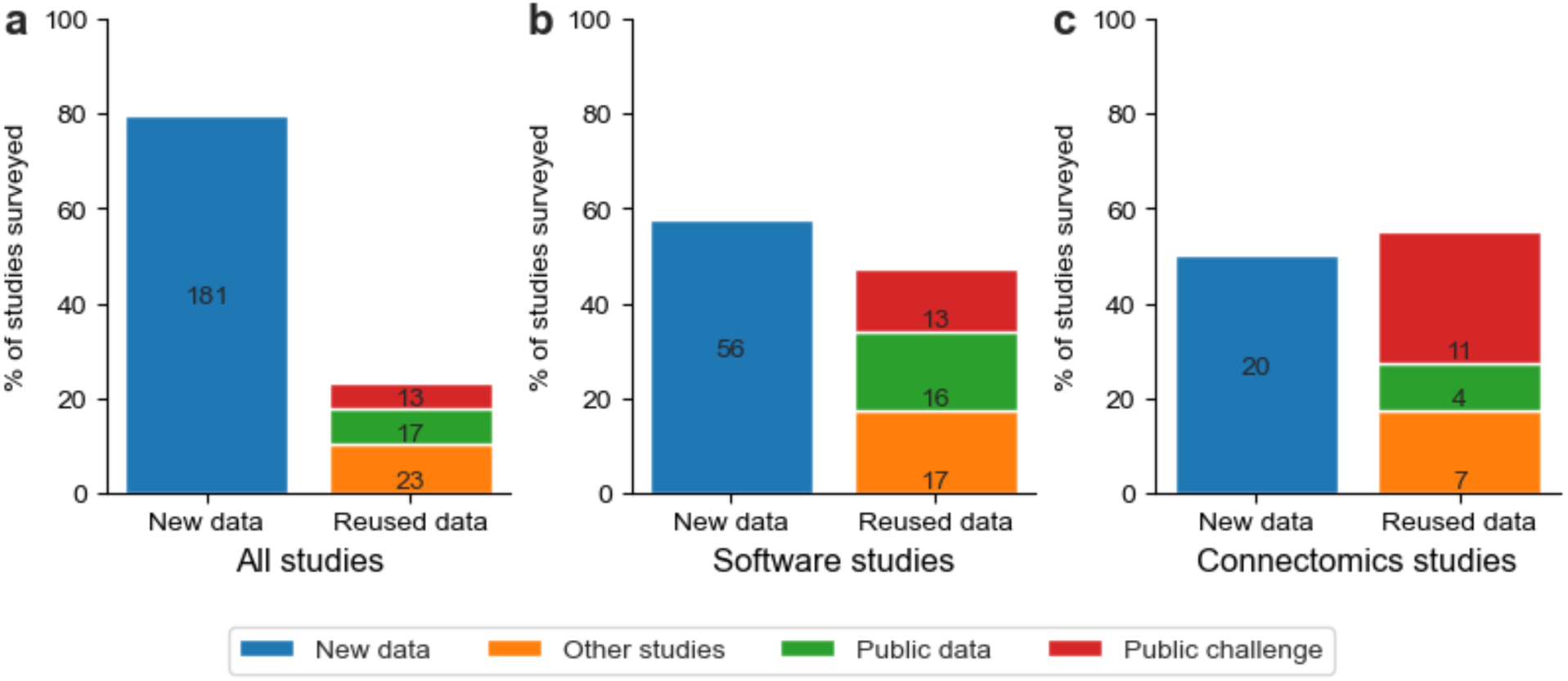
Sources of data used in all publications surveyed (n=227), only publications describing a software tool (n=97), and publications in the connectomics field (n=40). The number of publications in each sub-category of data source is displayed above the bar. Most publications collected their own data, including when the purpose of the publication was the description of a software tool. The Connectomics field demonstrated the best rates of re-use with many publications referencing the data from a public challenge. Note that some papers may be counted multiple times, e.g., a paper describing a software tool used in connectomics would be counted as a software, connectomics, and in all papers.

Reused data was most often sourced from other studies, which may be less openly accessible than data from public repositories as often these studies were conducted by the same research group, which was the case for 20 of the 23 studies surveyed sourcing data from other studies. Public data was also commonly used, but public challenges had the most reuse per dataset. One notable example from the connectomics field is that 10 of the 40 total connectomics publications used data from the ISBI 2012 challenge (Arganda-Carreras, et al., 2015). However, this data is no longer available at the original website (a public challenge site), making it no longer findable (Note: it is still available at https://imagej.net/events/isbi-2012-segmentation-challenge).

### Differences between imaging modalities

X-ray micro-computed tomography publications had the lowest rate (5.9%) of data re-use (Figure 4). Room-temperature EM (FIB-SEM, SBF-SEM, EM, ET, ssTEM, ATUM SEM, room-temperature ET, ssSEM) publications had the highest rate of reusing data collected elsewhere, with 35.8% of publications obtaining data from other studies, public data, or public challenges. This high rate of re-use is mainly due to the connectomics field. CryoEM (cryoEM, cryoET, cryoFIB-SEM) publications had a slightly lower rate of reusing data (31.4%).

**Figure 4.**
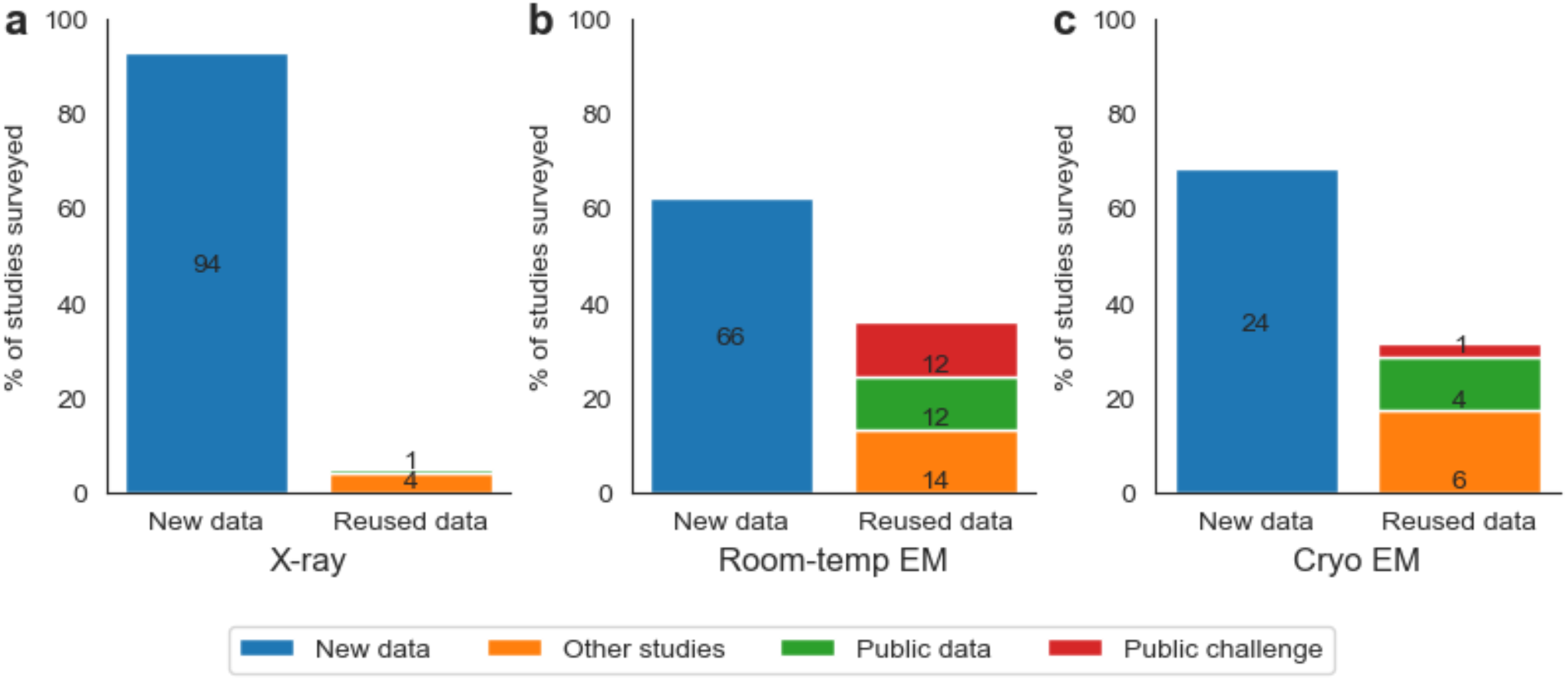
Rates of data reuse in publications from X-ray (n=101), room-temperature EM (n=94) and cryo-EM (n=31). Note that some publications were counted in several categories if multiple modalities were used.

Across all data types, cryoEM publications were most likely to deposit data in findable and accessible locations such as field-specific databases, general scientific databases or self-maintained websites (Table 1). Room-temperature and cryoEM studies most frequently deposited their image data (image training, and label) into field-specific databases such as EMDB or EMPIAR (Supplementary Figures 2-4). The X-ray publications with image data depositions were evenly spread across field-specific databases, general scientific databases, and self-maintained websites (Supplementary Figure 2).

**Table 1.**
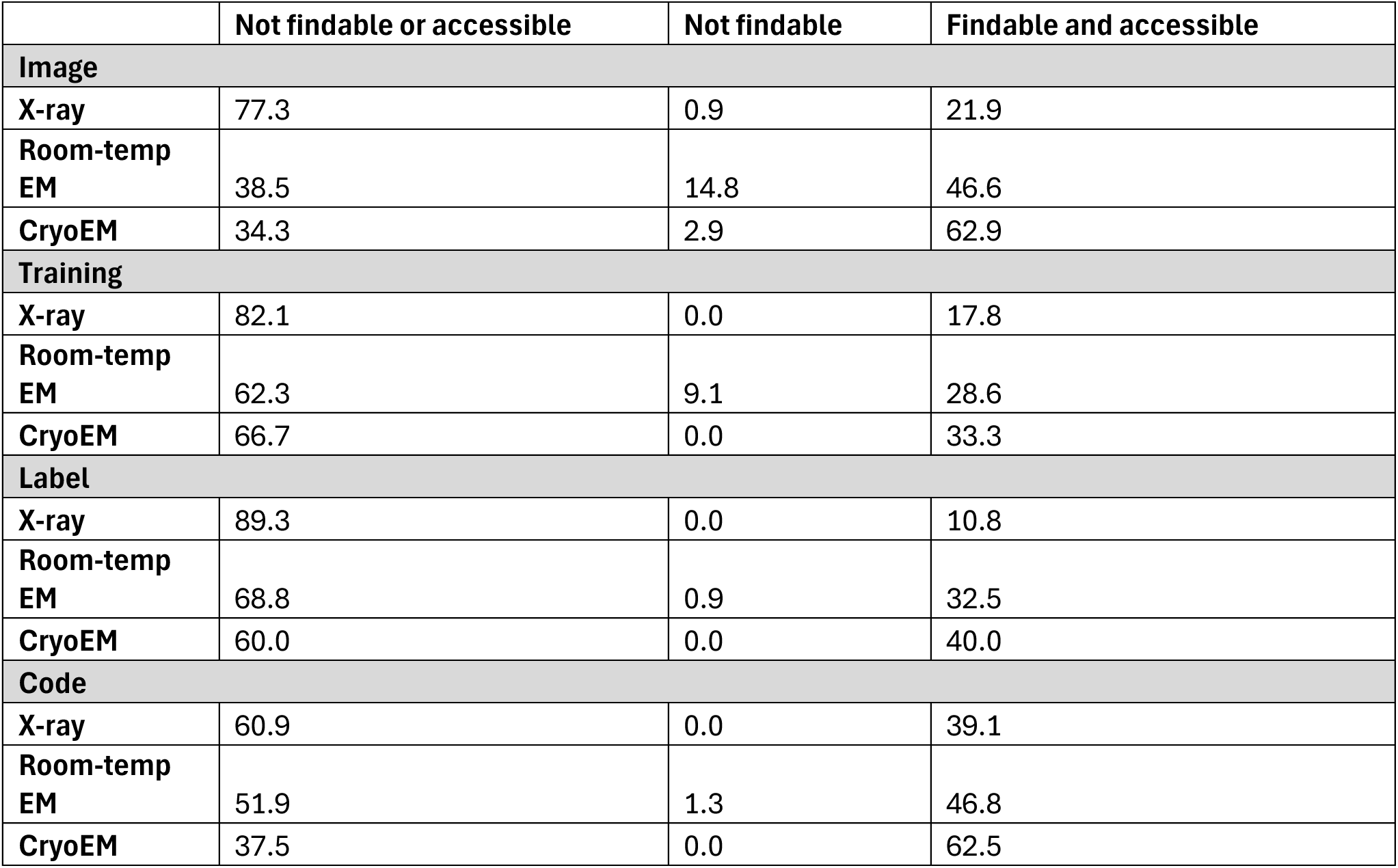
Percentage of publications in X-ray (n=101), room-temperature EM (n=94) and cryo-EM (n=31) depositing diDerent types of data in each type of location.

## Discussion

### Ontological considerations

Many publications we reviewed were not findable or accessible as they were deposited at defunct URLs, were only available on request, or required researchers to search separately through citations or online to download the data. We found that data was deposited across at least 26 different file formats, and 32 different online repositories, limiting the interoperability of these datasets as significant effort would be required to parse these formats into a single unified database for training segmentation tools. Once found and accessed, we determined that there was considerable variation in the definitions of certain terms such as “reconstruction”, “mask”, “segmentation”, and “ground truth”. A consensus or clearer descriptions are required across the breadth of the bioimaging community to ensure that these datasets can be re-used appropriately without misinterpretation. Some recommendations on clearer descriptors of common bioimage data are included in Figure 5.

**Figure 5.**
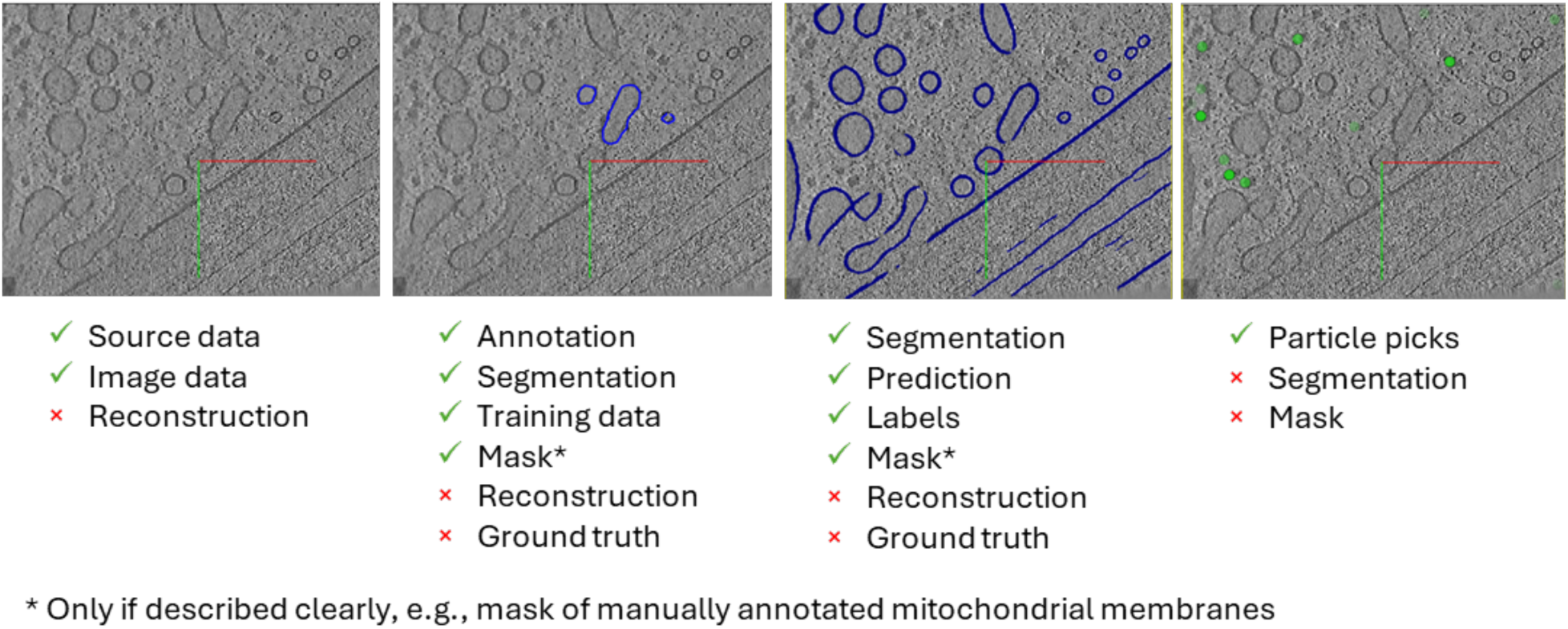
DiDerent common types of bioimage data and recommendations on terms that should and should not be used for clear descriptions.

The term ‘reconstruction’ is used often in bioimage research. It is colloquially used to mean any of the following:

- A 3D volume created from 2D images
- An average of multiple 2D sub-images or 3D sub-volumes into a 3D representation of a feature of interest
- A segmentation from a 2D or 3D volume

These various uses of the term ‘reconstruction’ lead to confusion, and arguably none are needed, as the better descriptions for the above examples would be tomogram/3D image data, averaged feature and segmentation respectively.

The term ‘mask’ can similarly cause confusion as it is used to describe partial annotations created as training data, predictions from semi- or automated approaches and proof-read outputs. To a non-expert, each of these binary images may be present with little or no associated description in a single repository entry. There may even be multiple labels present in the same file with no associated metadata to describe the objects being labelled; or when these descriptions are present, they are often shorthand or non-standardised vocabulary (e.g., “mito” or “endo” to describe mitochondria or endosomes respectively).

The term ‘segmentation’ is canonically used to describe the pixel- or voxel-wise labelling of features within an image, but recently in the cryoET field has also begun to be used to describe the practice of re-inserting an object’s average back into a 3D volume based on the orientations and locations where individual objects were found prior to averaging. This new practice is subtly different in a meaningful way, especially to non-experts, as the re-inserted averaged object may not actually be present at that location, may not be found in the orientation it is displayed in and may not represent the actual density present in the image(s). A more appropriate term for this use-case would be averaged re-insertion annotation (described by a set of coordinates and a vector) to distinguish between original particle annotation. In this way, the object’s locations and orientations can be provided alongside of the single average that represents all members of the group, reducing the computational storage burden. Importantly, we are not arguing that the practice of re-inserting averaged objects be stopped, just that it is better described and differentiated from the canonical use of the term ‘segmentation’.

The issue of ground truth is more insidious; in the bioimaging field, it is often used to describe expert annotation/segmentation. However, the term ground truth indicates that it is infallibly accurate because the truth is known (e.g., from a simulation). When the term “ground truth” is used to describe imperfect, biased, manual annotation/segmentation it can lead to interpretation issues where experts disagree or where automated/semi-automated strategies are erroneously discounted because they differ from the expert annotation (Hecksel, et al., 2016). The best term to use here is either manual annotation/segmentation or expert annotation/segmentation.

Volumetric imaging is often a long process of sample preparation, data collection and data processing/analysis. Choosing to use careful language when describing this process in publications, repositories and code gives the best chance of both shared understanding and re-use, especially beyond the immediate research field.

### The need for metadata

Segmentation specific metadata has lagged behind metadata associated with image acquisition, but it is crucially important to capture metadata from what is often the last step of the process, where the most energy has been expended, to reduce this burden in future. On a basic level, information as simplistic as what the feature of interest is (ideally linked to standard tissue and cellular ontologies) and whether it is described by zeros or ones (or something else) in the associated segmentation file would be a great place to start.

A secondary step would be to capture metadata associated with the purpose and quality of a segmentation and the process by which a segmentation was generated. There will likely be quality differences between segmentations generated for exclusively visualisation purposes versus those generated for quantitative analysis. And it is also likely that those segmentations which have undergone proofreading post-automated segmentation or by a second expert will be of higher quality than those which have not.

Beyond the basics, enhanced, automated metadata capture and search capabilities would help developers find suitable datasets for their use case and determine how much (if any) manual curation would be required to bring segmentation data to the required standard for their study. This metadata could include the number and size of the feature being segmented relative to the image, or whether the segmentation was verified by a correlative imaging approach (e.g., fluorescence to confirm location of a feature of interest in EM).

### Specific recommendations

Action is needed at different levels to work towards FAIR deposition of segmentation data in the bioimaging community (Figure 6). This has been previously understood in the context of capturing the details and data associated with individual imaging modalities (EMDB, EMPIAR, BioImage Archive, IDR), and recommendations have been made around minimum metadata standards across entire experiments and between imaging modalities (e.g, REMBI (Sarkans, et al., 2021)). However, segmentation is possible across many different individual imaging modalities with direct links to biological ontologies and non-imaging experts, meaning the mixture of practices is particularly broad and is not yet addressed by an individual modality or set of recommendations. Many of the recommendations made here are also present in the recommendations/standards for individual modalities; those that are more specific to segmentation are common sense for data/metadata capture and reuse. In all cases, these have been discussed with relevant communities and presented for discussion in national and international forums. Action is needed at different levels to work towards FAIR deposition of segmentation data in the bioimaging community (Figure 6).

**Figure 6.**
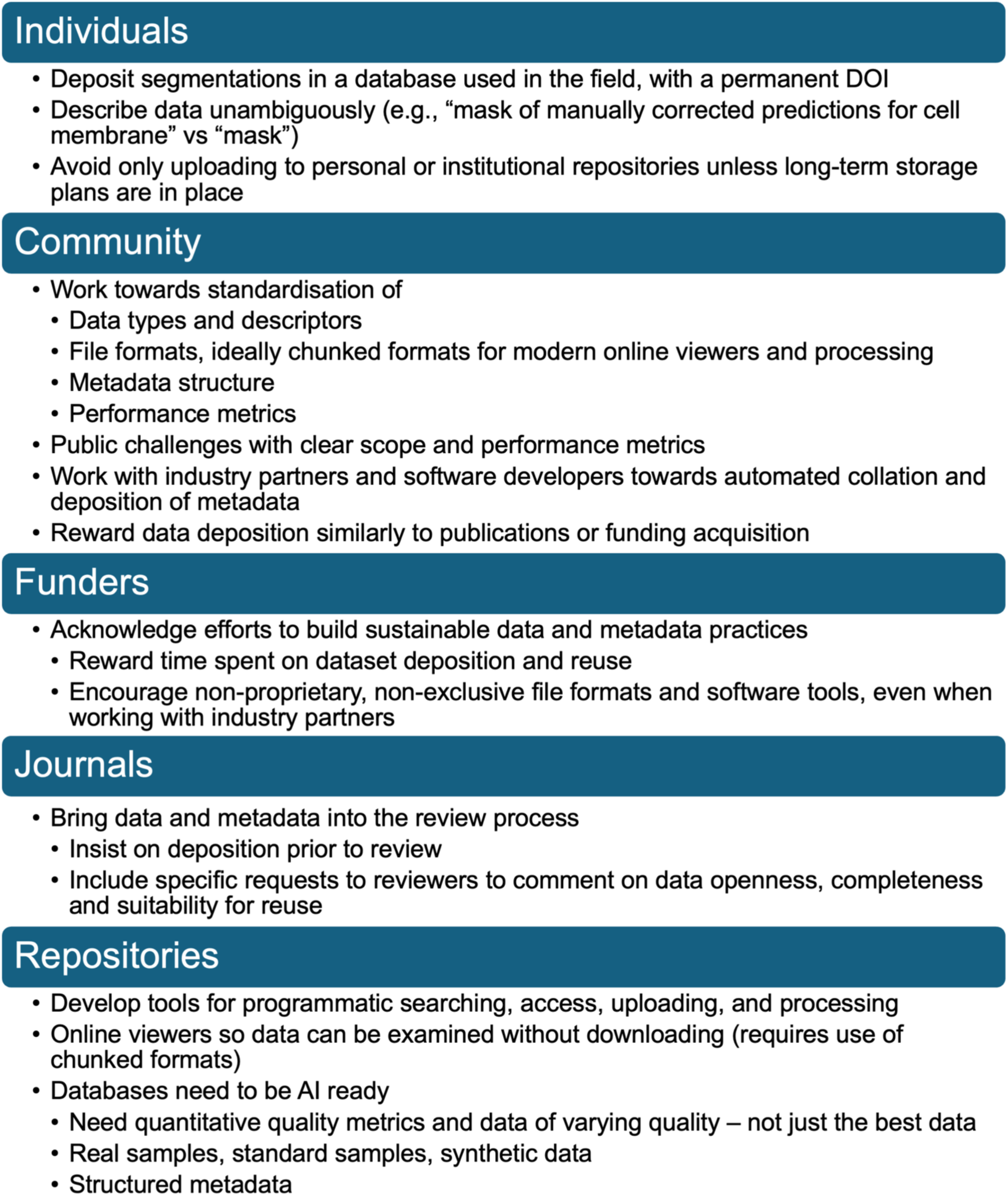
Specific recommendations for action at diDerent levels to improve FAIR deposition of segmentation data in bioimaging.

As an individual researcher please deposit your data, training data, label data and code. This includes segmentations that were done for visualisation purposes (e.g., show that subcellular organelles were visible in the images), and especially segmentations which formed the basis of quantitative conclusions (e.g., count and measure volume of mitochondria in images). Choose a field-specific database that provides a permanent DOI for your data; avoid institutional or personal repositories or those which are not permanent. Using field-specific databases allows concentration of community efforts in developing the computing infrastructure to store, index, and curate data which can address field-specific concerns, e.g., EMDB including automatic quality reporting for electron density maps of protein structures. It is much more difficult to develop a ‘one-size-fits-all’ framework for all types of microscopy, especially as the field explores correlative experiments, so general databases such as the BioImage Archive or Image Data Resource are invaluable in hosting those datasets before they become common enough to require more bespoke solutions tailored to the specificities of the field. When you are writing your manuscript and depositing your data, describe it clearly and unambiguously – consider the next researcher who will be tasked with attempting to reuse your data. When reviewing manuscripts, insist on the data, training data, label data and code being deposited in a field-specific database before acceptance.

As a community, we must standardise the data types and descriptions, file formats and metadata associated with our data, training data and label data. This is hard, but it is important. We must also move towards chunked file formats (e.g., OME-Zarr (Moore, et al., 2023)) to allow for online viewing and processing of large data. Unsurprisingly, public challenges play a large and unique role to move the field forward; they should be encouraged and new performance metrics should be developed to understand the outputs. Care should be taken to ensure the metrics used during challenges align with desired outcomes. We must work with our industry partners and software developers to ensure appropriate metadata is generated instead of black-box tools, and to automate the collation and eventual deposition of this metadata. As a community we must also reward deposition of data similarly to how we reward manuscript publication or funding acquisition.

As a funder, please acknowledge the efforts of individuals and communities to build more sustainable data and metadata practices. Please consider dataset deposition and reuse as valid outputs and reward time spent towards these endeavours. Some steps are being taken in these directions by some funders (e.g., UKRI’s recent data policy draft document). Please also enable collaborative industry research while ensuring that public funds used to support these collaborations are not used exclusively to further industry partner market position by encouraging non-proprietary, non-exclusive adoption plans.

As a journal, please bring the data and metadata into the review process by insisting on deposition prior to review and including specific requests to reviewers to comment on the openness and completeness of the deposited data; it’s suitability for reuse. This will enable the data to be assessed more easily during the review process and will ensure it is already deposited prior to publication. This is common practice in some structural biology fields (e.g., macromolecular crystallography).

As a repository, please provide tools for programmatic searching, access, uploading, and processing of data, including the adoption of chunked file formats. This will enable developers and researchers to understand the data before expending the effort and resources to transfer a copy of it to local storage, or better yet, alleviate this need entirely. And finally, we need to consider AI within the context of volumetric imaging and get our data ready. We should ensure there is a place for standard samples and synthetic data alongside of the “real” data. This includes understanding the quality of the data using quantitative metrics and the deposition of data of variable quality – not just the best data.

## Conclusion

Data deposition rates are disturbingly low, especially data from steps beyond the initial imaging data. Re-use of data is essential for many purposes, not least the development of segmentation software tools, particularly those using artificial intelligence techniques requiring large amounts of training data. Clear, unambiguous descriptions of data and metadata will democratise the development of solutions, broadening beyond the original field and scientists associated with large monetary investments in microscopes and associated kit. Changes are needed at all levels from individual researchers, the bioimaging community, and the repositories themselves; but together we can ensure precious data is captured for re-use and we are prepared to take advantage of future advancements in AI.

## Methods

A collection of publicly available segmentation and annotation datasets for 3-D volumetric electron microscopy and associated metadata was assembled from searches in the EMDB, EMPIAR, and Open Organelle databases, and a literature search with Pubmed from 2012-2023. Characteristics about these datasets were collected and trends over time were investigated, e.g., the purpose of the segmentations, data formats, biological feature type and size scale, imaging modality, and others.

### Literature searches

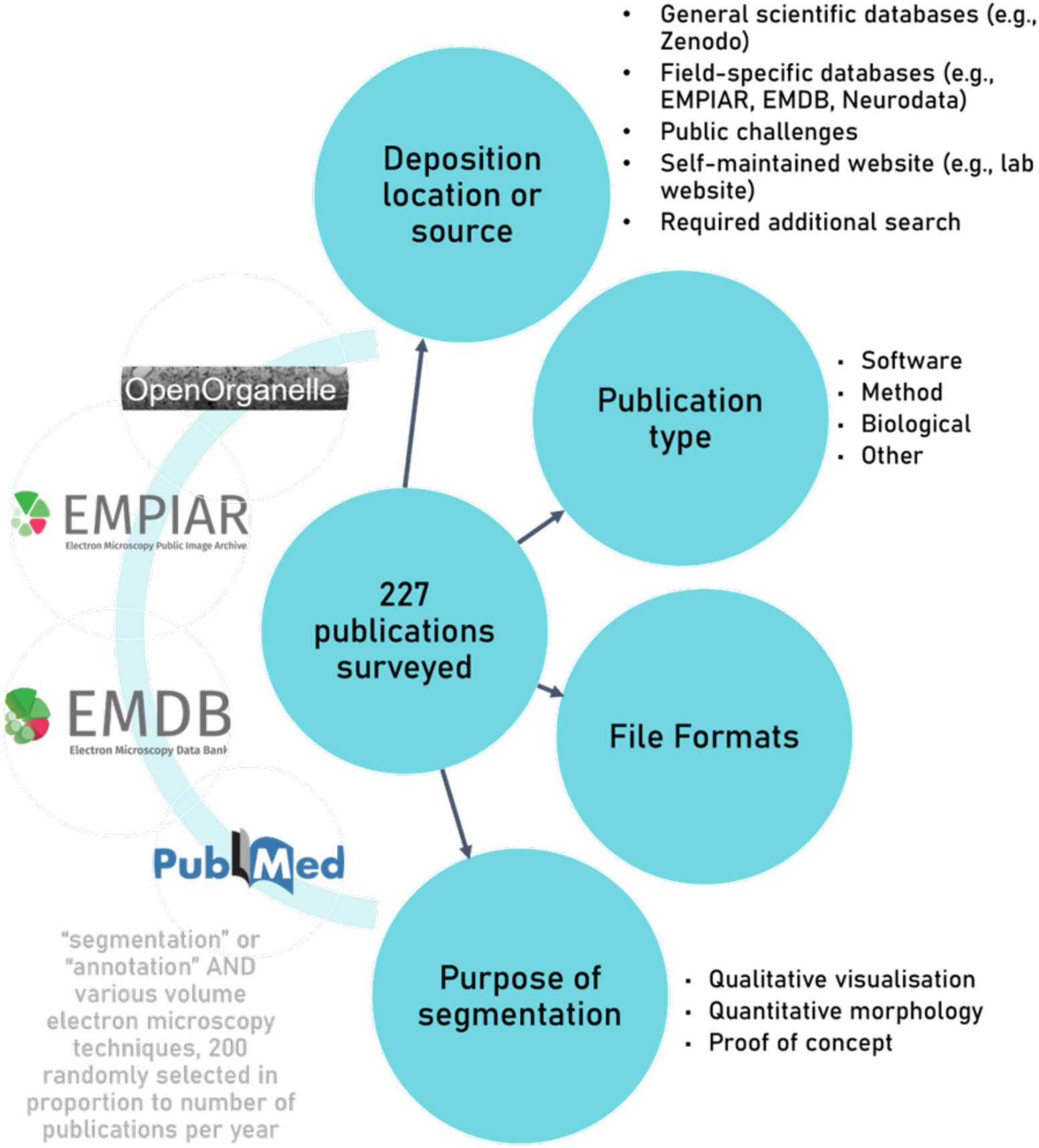

Searches were conducted on Pubmed and aggregated. Duplicates were removed and 100 publications were selected randomly from the remaining 2125 publications, in proportion to the relative number of publications for each year from 2014-2024. This sampling strategy meant that publication years with fewer overall publications matching the search terms had fewer publications added to the final sample ensuring representation of overall trends in the fields and across time were preserved. Selection was performed by assigning every publication a random ID number, the search results were then filtered by year of publication, and the publications were sorted in ascending order of the randomised ID. Before a paper was added to the sample pool, it was reviewed more closely to determine if the paper was relevant to the bioimaging community and questions at hand (i.e., delineation of biological features of interest from an electron microscopy image). This was repeated until the total number of publications for each year were added to the sample pool to reflect the proportion of publications matching the search terms per year.

The specific searches used to represent the volume electron microscopy field were:

- “((“segmentation”) OR (“annotation”)) AND ((SBF SEM) OR (SBEM) OR (serial block face scanning electron microscopy))”
- “((“segmentation”) OR (“annotation”)) AND ((FIB SEM) OR (FIB-SEM) OR (FIBSEM) OR (focused ion beam scanning electron microscopy))”
- “((segmentation) OR (annotation)) AND ((electron microscopy) OR (EM))”
- “((“segmentation”) OR (“annotation”)) AND (electron tomography)”
- “((“segmentation”) OR (“annotation”)) AND ((ssTEM) OR (serial section TEM))”

The Pubmed search was repeated for the X-ray imaging field and 100 publications were selected randomly from the 5651 publications, in proportion to the relative number of publications for each year from 2014-2024 (as described above). The specific search used to represent the X-ray imaging field was:

- “((“segmentation”) OR (“annotation”)) AND ((X-ray microscopy) OR (X-ray microCT) OR (X-ray tomography) OR (X-ray micro-computed tomography))”

EMDB and EMPIAR were searched using the following query:

- “segmentation_filename:[* TO *] AND structure_determination_method:tomography”, “segmentation”, “segmentations”, “annotation”, and “annotations”.

The OpenOrganelle database was manually searched for datasets that included image data and an annotation or segmentation.

The resultant search results from EMPIAR, EMDB, and OpenOrganelle were added to those results from the PubMed search and duplicates were again removed based on DOI. This enriched the sample pool ensuring questions related to current trends/habits of data deposition could be captured. The results were organised by publication, i.e. each publication was a single record regardless of how many datasets were uploaded. The 231 publications were reviewed and information about the datasets generated from the publications were recorded. These characteristics included:

- the study type (biological investigation, new imaging or sample preparation method, dataset, or software paper)
- segmentation use (proof-of-concept, qualitative visualisation, quantitative morphometry, single-particle analysis or sub-tomogram averaging)
- where the image data was deposited to (if at all)
- where training data was deposited to (if at all, or required)
- where label data was deposited to (if at all)
- where code for generating the segmentations was deposited to (if at all, or required)
- imaging modality
- file formats
- source of the datasets (collected for that publication, reused from elsewhere, publicly available)
- Segmentation type (semantic or instance)
- Segmentation method (automated, semi-automated or manual)
- Biological scale (tissue, cellular, sub-cellular)

## Data and Code Availability Statement

The spreadsheet containing the search results and code used to generate the plots are available at (10.5281/zenodo.14587760) or https://github.com/rosalindfranklininstitute/FAIRly-depositing-data.

## Author Contribution Statement

MCD conceived of and supervised the project. EMLH and MCD defined the search strategy and parameters. EMLH and DL collected the data. EMLH performed the analysis and prepared the figures. EMLH and MCD wrote the manuscript. All authors reviewed the manuscript prior to submission.

## Acknowledgements

The authors wish to acknowledge Wellcome Trust funding (Electrifying Life Sciences 220526/Z/20/Z)

## Supplementary Information

**Supplemental Figure 1.**
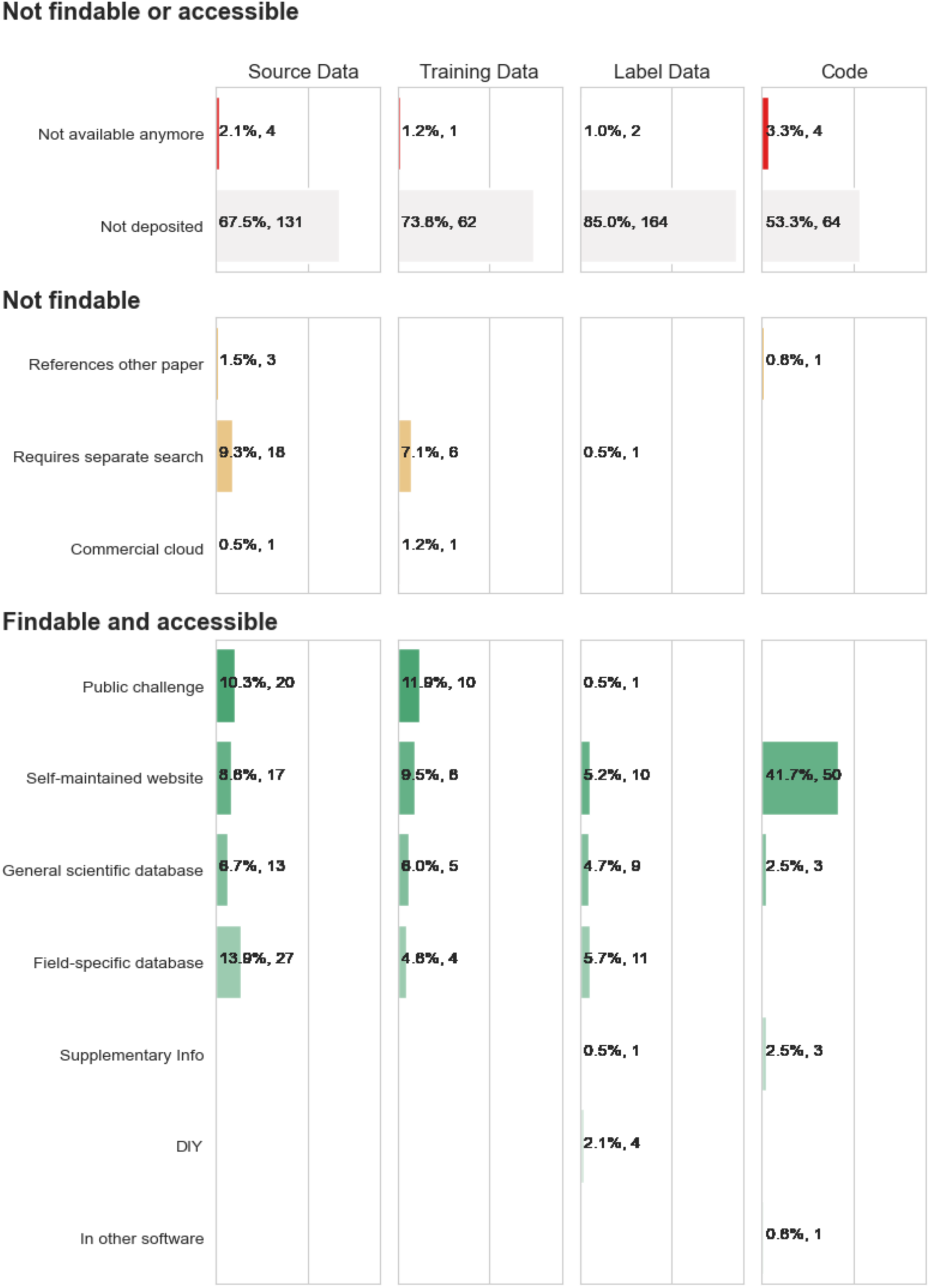
Deposition locations of data only from Pubmed searches, excluding those from the field-specific database searches.

**Supplemental Figure 2.**
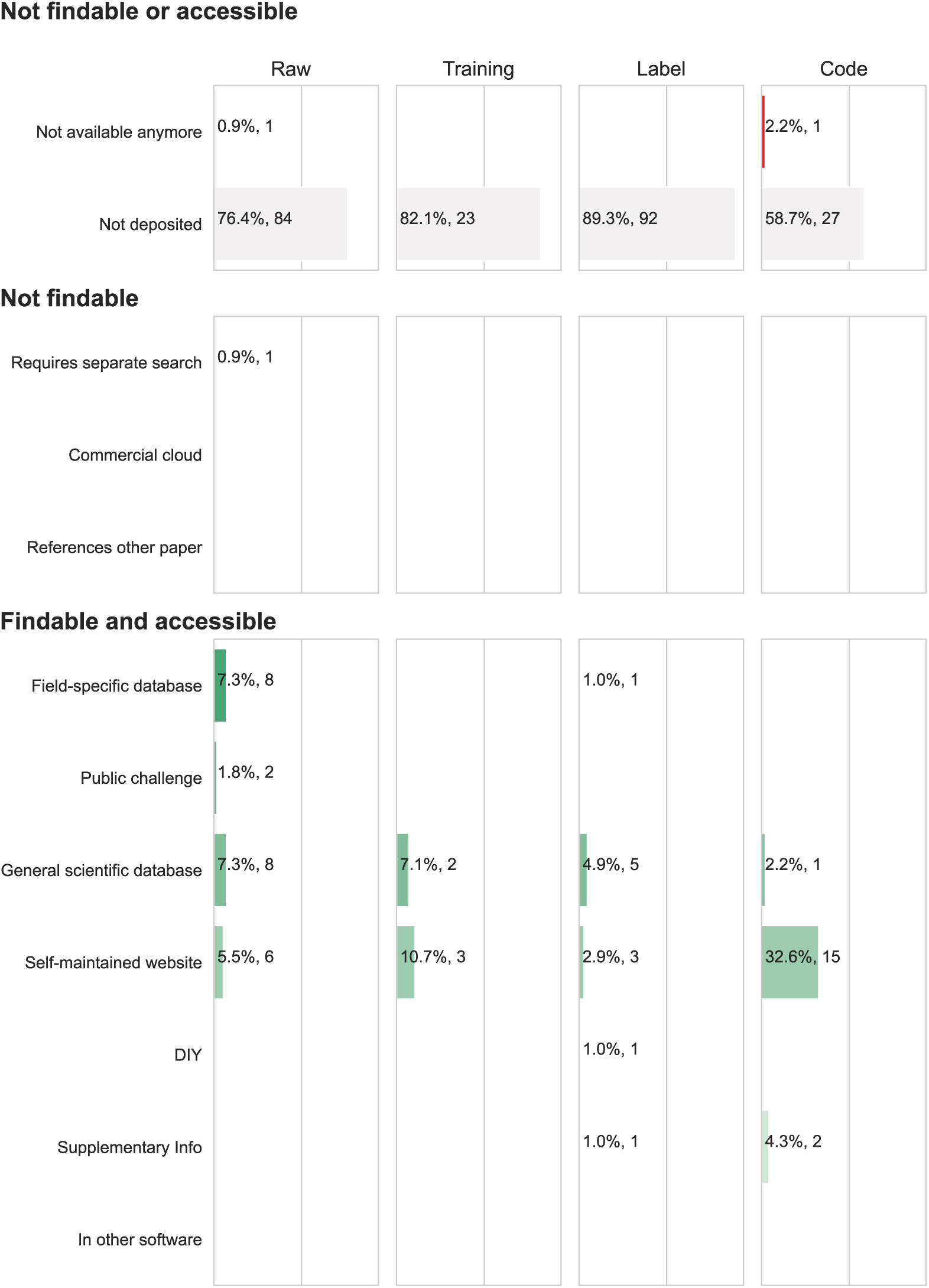
Deposition locations of image training, label data and code for X-ray publications (n=101).

**Supplemental Figure 3.**
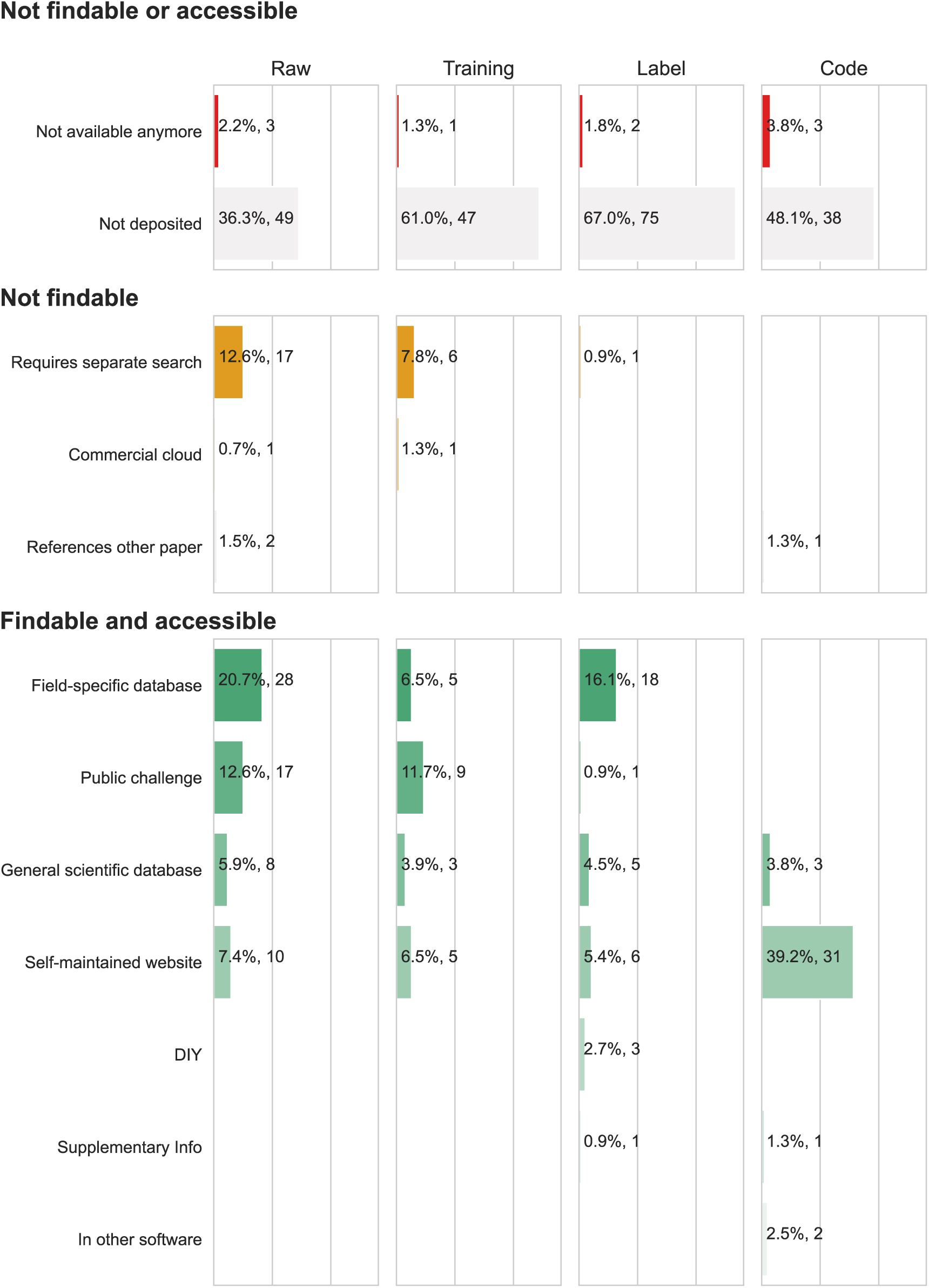
Deposition locations of image training, label data and code for room-temperature EM publications (n=94).

**Supplemental Figure 4.**
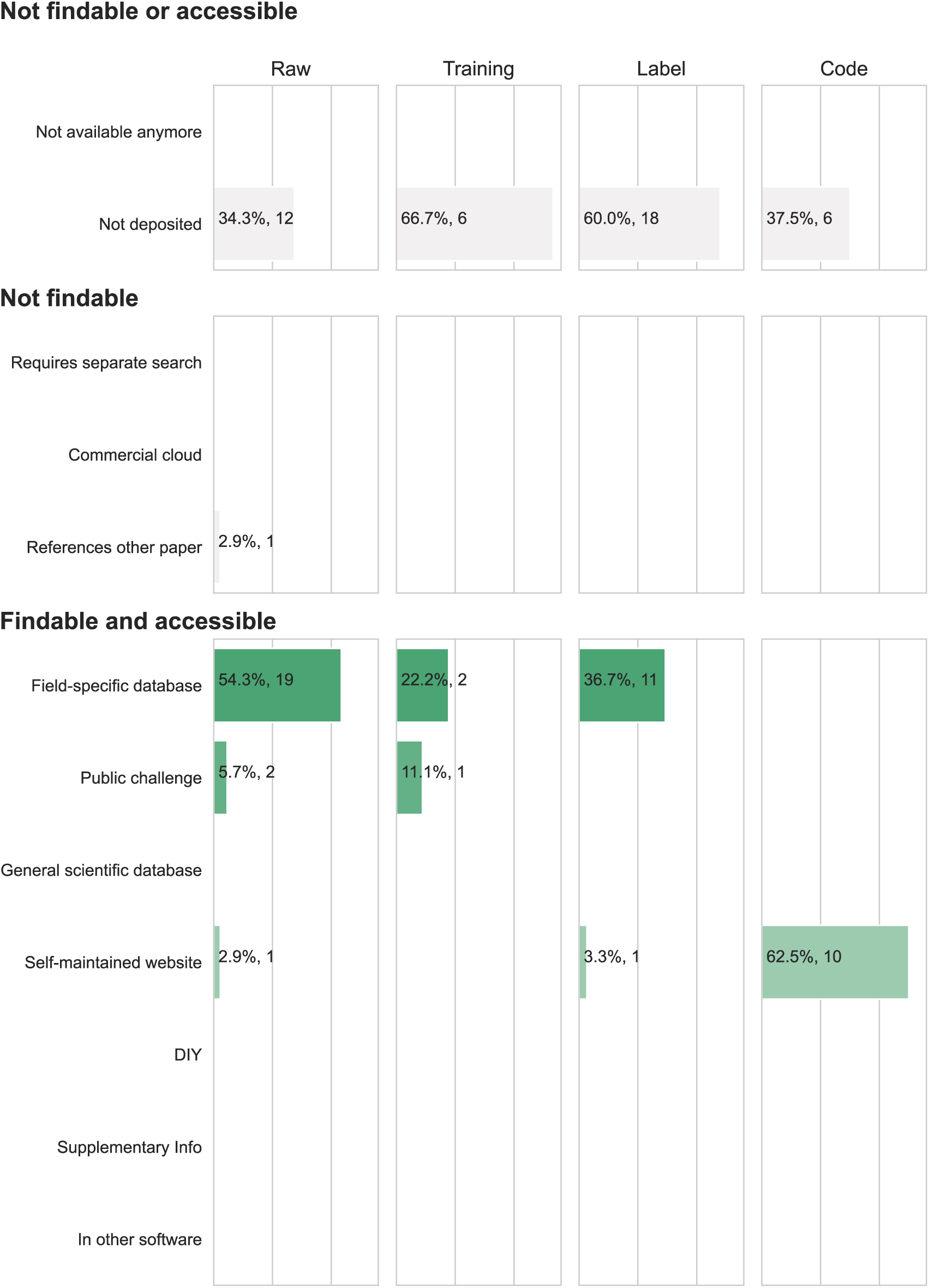
Deposition locations of image training, label data and code for cryo-EM publications (n=31).

